# Maximum entropy networks predict fluctuations and stability of food web energetics

**DOI:** 10.64898/2026.01.19.700332

**Authors:** G. V. Clemente, T. Caruso, M. Chomel, J. M. Lavallee, F. T. de Vries, M. D. Bustamante, M. Emmerson, D. Johnson, R. D. Bardgett, D. Garlaschelli

## Abstract

A central goal of ecology is understanding how the architecture of food webs, which represent the structural backbone of ecosystems, affects their stability. The analysis of stability in the classical sense of population dynamics (i.e. return to equilibrium) can be successful for a single instance of an empirical food web but ignores the multiplicity of alternative states in which the system could be found as a result of intrinsic variability and fluctuations. Here we propose and test a new methodology to reconstruct, from single empirical observations of a food web, the viable ensemble of alternative realizations respecting the observed resource-consumer linkages and empirical energetics. The reconstruction can be handled analytically within a maximum-entropy framework which predicts how empirical food webs access a multitude of alternative states with comparable stability and reactivity. The (measurable) entropy of the reconstructed ensemble directly quantifies this multiplicity and serves as a novel proxy of system resilience, that is the rate of return to equilibrium in response to an external perturbation. We show that the associated ensemble fluctuations provide explicit predictions for the expected response of food webs to external perturbations, such as anthropogenic or climate-induced stresses. We do that by validating the proposed fluctuation–response relation on empirical soil food webs subjected to experimentally controlled perturbations, confirming that intrinsic fluctuations in the unperturbed state predict responses to subsequent stresses. The perturbed states are associated with higher entropy, indicating less likely spontaneous recovery.

## I. MAIN

The diversity-stability debate [1–4] is a significant example of the extensive research efforts aimed at understanding how the structure of complex ecological systems influences their dynamics, particularly in response to perturbations. A central challenge in this field is integrating the substantial volume of theoretical and empirical studies, given that many theoretical frameworks and predictions remain difficult to experimentally validate or test [5–8]. One specific difficulty arises from the multiple, often multidimensional definitions of “diversity” and “stability” employed across studies [5, 9, 10]. Many theoretical and experimental ecologists have tackled the broad topic of the stability of complex systems by focusing on the major study case of food webs, examining how complexity and diversity affect their stability. A significant portion of this research leverages the mathematical properties of systems described by coupled nonlinear differential equations. These equations represent species as nodes in a network and quantify how each species impacts and is impacted by others through consumer-resource interactions [2, 3]. Complementary approaches have focused on analyzing the structural properties of food webs, attempting to identify links between network configuration and ecological stability [11–13]. Despite significant advancements, a critical knowledge gap persists between theoretical developments and empirical parameterizations required to test these theories. Bridging this gap has been partly achieved by research linking energetic descriptions of food webs with classical population dynamics [8, 14, 15]. For example, theory and some experimental evidence suggests that asymmetry of energy channels coupled by top predators is an important structural element that stabilise the population dynamics underpinning soil food webs [16].Other more theoretical works have offered criteria that dynamical systems should fulfil to be stable [1], with some empirical evidence supporting, if not proving, those criteria [17]. There is, however, no or very little theory and experimental work that links fluctuations to the factors and structural properties hypothesised to control the stability of food webs and their response to perturbations. Yet, fluctuations are one of the most important characteristics of natural systems given the complexity of the very many factors that simultaneously affect natural populations. For example, global change implies the simultaneous actions of many perturbing factors and increased spatial and temporal variance of natural system, which makes the modelling of fluctuations essential if we are to predict responses to future perturbations [18, 19]. We propose that the application of statistical physics theory to the description of network systems [20–22] offers the tools to simultaneously describe fluctuations in the energy fluxes and link these fluctuations to the stability of the food web, and its future response to perturbations. We demonstrate this approach theoretically and then apply the theory to an experimental soil food web, involving 30 pairs of control-perturbed grassland soil systems observed over two months following a perturbation regime [23].

Theoretically, we unify maximum entropy network models and energetic food web models through three steps: modeling food web energetics as networks described by energy flux matrices [14, 15], defining entropy as the number of typical equilibrium configurations [22], and employing the fluctuation-response relation applied to these networks [24–26]. All three steps can be applied to experimental data, as shown below.

First, we follow the procedure described in Box I, applying the maximum entropy principle to identify the probability distribution that is maximally unbiased while still reproducing a prescribed set of constraints. Among the various models based on different types of constraints [18, 22, 28], in this work we adopt the *Conditional Reconstruction Method* (CReM) [29], specifically its “CReMa” variant (Supporting Information, Sec. 1). In this formulation, the network topology is fixed by the adjacency matrix, while fluxes are randomized under the condition that inflows and outflows of each trophic species are preserved *on average*, that is, the constraints are satisfied only in expectation, thus allowing for fluctuations. Within this framework, we can incorporate the experimentally observed flux matrix directly into the graph Hamiltonian of the CReMa model (SI Sec. 1), which then becomes

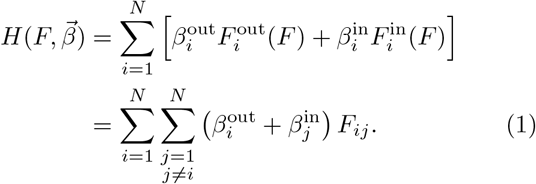

Where **F** is a weighted matrix of energy fluxes linking the species in the food web. For the experimental derivation of the energy flux matrix, we used [14, 15], with details summarised in the Methods (Sec.II B). The parameters can be estimated by maximum likelihood with some notable results (see Box I and Supporting Information 1) because the model can be resolved easily to find the probability of observing a **F** given the topology of the food web and the Lagrange multipliers *β* in the Hamiltonian. This formalism thus allows describe any energy based food web as a maximum entropy network in which local constraints are enforced only on average, that is a canonical ensemble in statistical physics [18, 22]. From here, we proceed to the second step, described in Eq. I.5 of Box I. This step allows us, after estimating the Lagrange multipliers, to compute the system’s entropy, which quantifies the number of typical configurations of the system [22]. In statistical mechanics, such ‘typical configurations’ correspond to the microstates that the system predominantly explores at equilibrium. Within our framework, this formalism can be exploited to interpret the number of typical flux configurations (or matrices) as the number of systems capable of reproducing a similar biomass *B*_*i*_ dynamics for each species. This depends on the general form of the dynamical system, but as we show, it is valid in our case and can, in principle, be extended to other cases. The link between the dynamics of and flux through a species, as described in Box III, is given by the mass balance equations [30]:

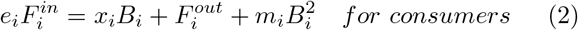

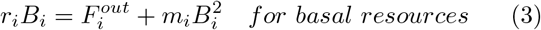

### Box I: From a single observed food web to an ensemble of alternative networks

**Step (a): Shannon–Gibbs entropy and constraints**. We define the entropy of a probability distribution *P*(*G*) over graphs *G* as

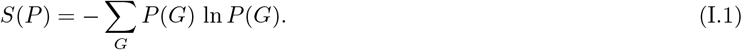

Prior knowledge is encoded by constraints on observables *C*_*i*_(*G*):

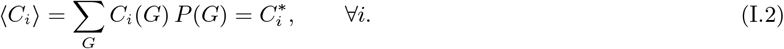

**Step (b): Lagrange multipliers and canonical ensemble**

Maximization of entropy subject to the above constraints yields the Boltzmann distribution [20, 21]

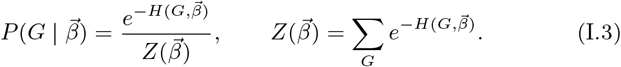

This defines a canonical ensemble of graphs [20], with 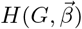 playing the role of the Hamiltonian.

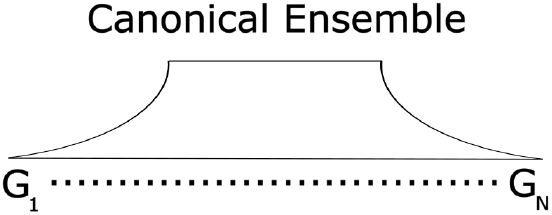

Canonical ensemble spanning accessible configurations.

**Step (c): Parameter estimation via maximum likelihood**. Given an observed network *G*^∗^, the parameters 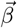 are determined by maximizing the log-likelihood [27]:

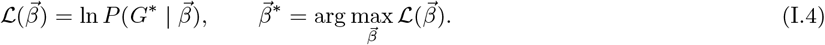

At the maximum, the log-likelihood equals the entropy of the model evaluated on *G*^∗^ [22]:

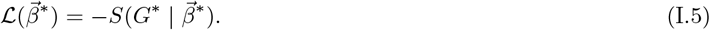

This procedure yields both (i) the fitted parameters 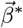 encoding the constraints, and (ii) the numerical value of the entropy consistent with the observed network *G*^∗^.

Where 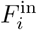 is the flux of energy entering species *i*, 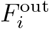 is the flux going out from species *i* to support its consumers; *e*_*i*_ is the feeding efficiency; *x*_*i*_ is the mass-specific metabolic rate and *B*_*i*_ the total biomass of species *i*; *m*_*i*_ captures the self-limitation strength and *r*_*i*_ is the growth rate for a basal resource.

We can connect the value of the entropy with the system dynamics: the greater the entropy, the greater the number of similar systems that reproduce a similar dynamics. This means that, given an average amount of total energy through the entire network, the network has multiple configurations in the distribution of the fluxes with the most typical one siting in the middle of the distribution.

As the canonical ensemble generated from the observed matrix of fluxes *F* ^∗^ can be sampled numerically to generate a desired number of randomised flux matrices, we have effectively derived multiple versions of the matrix flux that correspond to multiple dynamic systems. In other words, the randomization of the matrix *F* ^∗^ corresponds to that of the underlying dynamic systems, which solves the challenge of creating null models for food web dynamics.

For every flux matrix, the Jacobian of the corresponding population dynamics can be derived (see Methods Sec. II D and Supporting Information 5) to estimate classic metrics of resilience and resistance (Supplementary Information 4) of the systems [2, 30–33], which unifies a network description based on the energetic food web to a classic population dynamics model of food webs.

Our empirical findings (Supplementary Information, Fig. 5a,b) show that the resulting network ensemble accurately estimates the maximum eigenvalue for each of the 60 analyzed soil food webs associated with the Jacobian and its Hermitian. Consequently, our network model reliably predicts resilience, i.e., the return time to equilibrium following perturbations, and the *reactivity*. These results thus provide robust empirical support for a key hypothesis previously proposed in the literature: systems’ responses to perturbations directly depend on the input and output of energy flows from individual nodes and the network’s topology with consumers and top predators lining the energy channels [16, 30]. In fact, the model is also consistent with a prediction of the observed coefficient of variation of biomass over time (Supplementary Information, Fig. 5c), although the ensemble tends to overestimate this variance. Validating this results is crucial for interpreting the relationship between food web dynamics and entropy. It establishes a connection between a property of the ensemble, i.e., the network model, and the underlying dynamical system. Furthermore, entropy was found to be well correlated with the coefficient of variation, the magnitude of the leading eigenvalue, and the system’s reactivity (Fig. 1a-c).

**FIG. 1:**
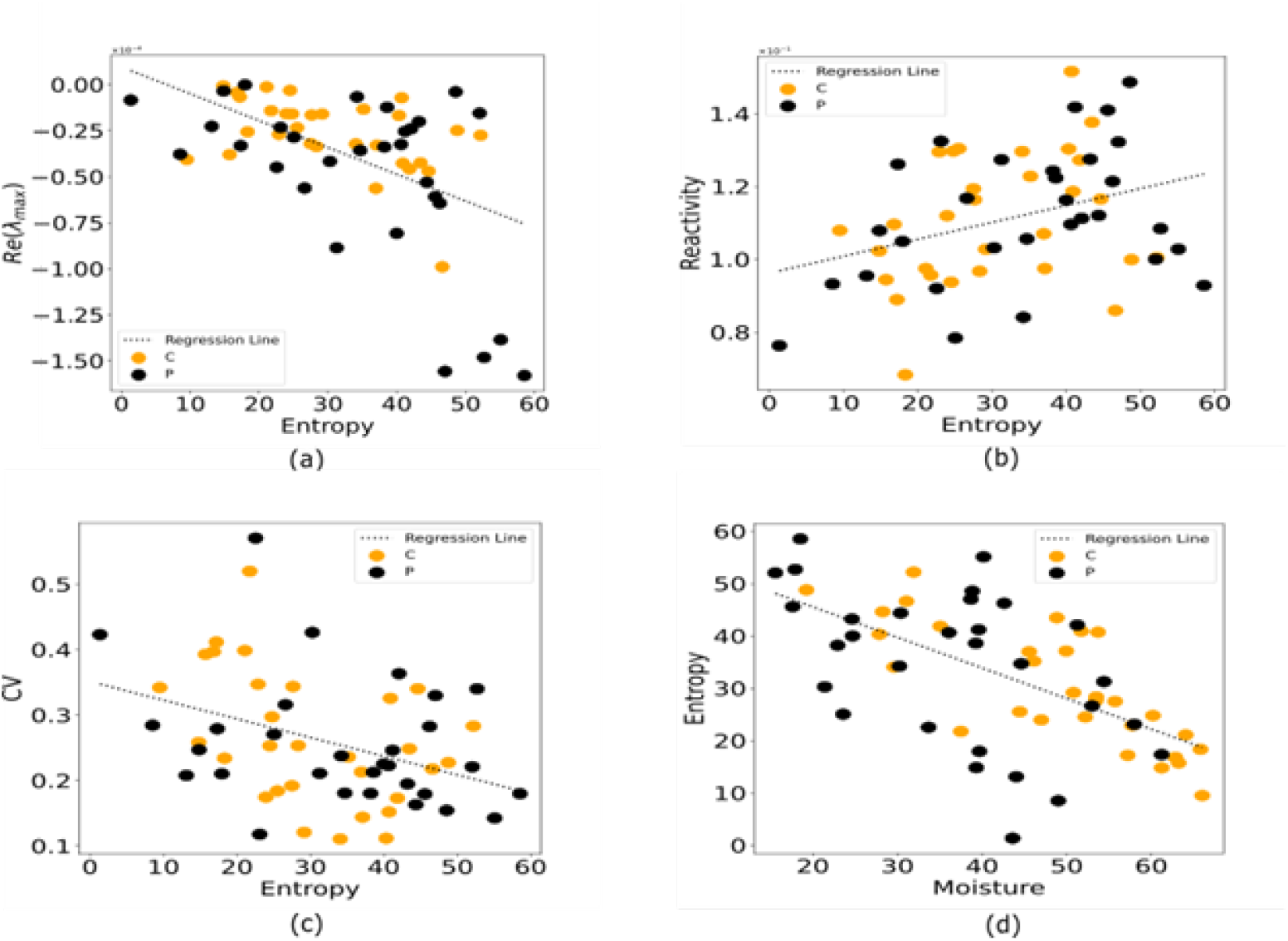
Scatter plot illustrating the correlation between the network entropy and the real part of the maximum eigenvalue (Re(*λ*_max_), panel a), Reactivity (panel b), the coefficient of variation (CV, panel c) and soil moisture (panel d), which is much reduced on drought food webs. Orange points (labelled *C*) represent control (unperturbed) systems, whereas black points (labelled *P*) represent systems subjected to drought.

A decreased moisture, which is caused by the perturbation of drought, is also associated with a higher entropy, which means a higher number of states visited by the system at equilibrium (Figure 1d). This higher number of states correlates positively with the magnitude of the real part of the maximum eigenvalue (which are negative), meaning increased resilience. That correlates also with a lower CV of biomass, meaning a more temporally stable biomass. Given the analytical relationship between the entropy and the total energy flow, a high energy flow implies, under similar conditions, a great capacity for network to reconfigure (i.e., a large number of compatible configurations). However, it is easy to show (Supporting Information 6) that entropy not only encompasses information about the total energy flow but also reflects how the total energy is distributed among its various components. Finally, in the third step, we derive the fluctuation response relation (See Box II) by considering the total flux through the food web *F*_tot_ with the goal of demonstrating that the knowledge of the value of *F*_tot_ before the perturbation allows predicting the new value of *F*_tot_ following the perturbation.

This choice to focus on a global constraint such as total flux is twofold: first, it stems from the established relationship between total flux and global system properties [35]; second, it reflects the global nature of the perturbations applied to the system, which justifies expecting the emergence of such a relation in global variables. This relation, applies to small perturbations and describes how fluctuations observed in a variable at equilibrium are connected to the system’s response to perturbations.

The relationship can be tested on our empirical data of food webs with paired control and perturbed systems. The data set consists of 30 pairs of control and perturbed food webs with both control and perturbed system observed four times after the release of the perturbation and in which the perturbation was experimentally imposed as a spell of drought [23]. To test the relationship, we considered that the perturbation acts globally on the entire system. For this reason, we used the variance of the total flux, computed from the model with the global constraint (see Section 2). The same result can also be derived from local constraints, as shown in the Supplementary Information (Section 3). Therefore, the total variance of *F*_tot_ at equilibrium takes the following form:

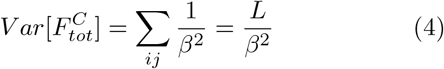

And defining:

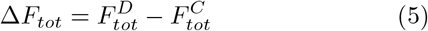

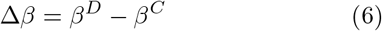

We can use these quantities to construct a proxy for the left-hand side of Eq. (II.4).

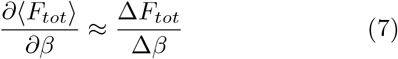

This models fit our date extremely well (Figure 2a) and it also allows connecting the variation in the structural variation of the network parameter beta (caused by the perturbations) to the intensity of the perturbation (in our case the difference in soil moisture between control and drought, the x-axis of Figure 2b). A greater perturbation (more negative Delta in moisture between control and drought) causes a greater variation in the network parameter. The greater the response of the fluxes to a perturbations are, the greater the fluctuations of the system at equilibrium are. Finally, the validation of the relationship under the assumption of a global effect of the perturbation can be regarded as evidence supporting the soundness of this assumption.

**FIG. 2:**
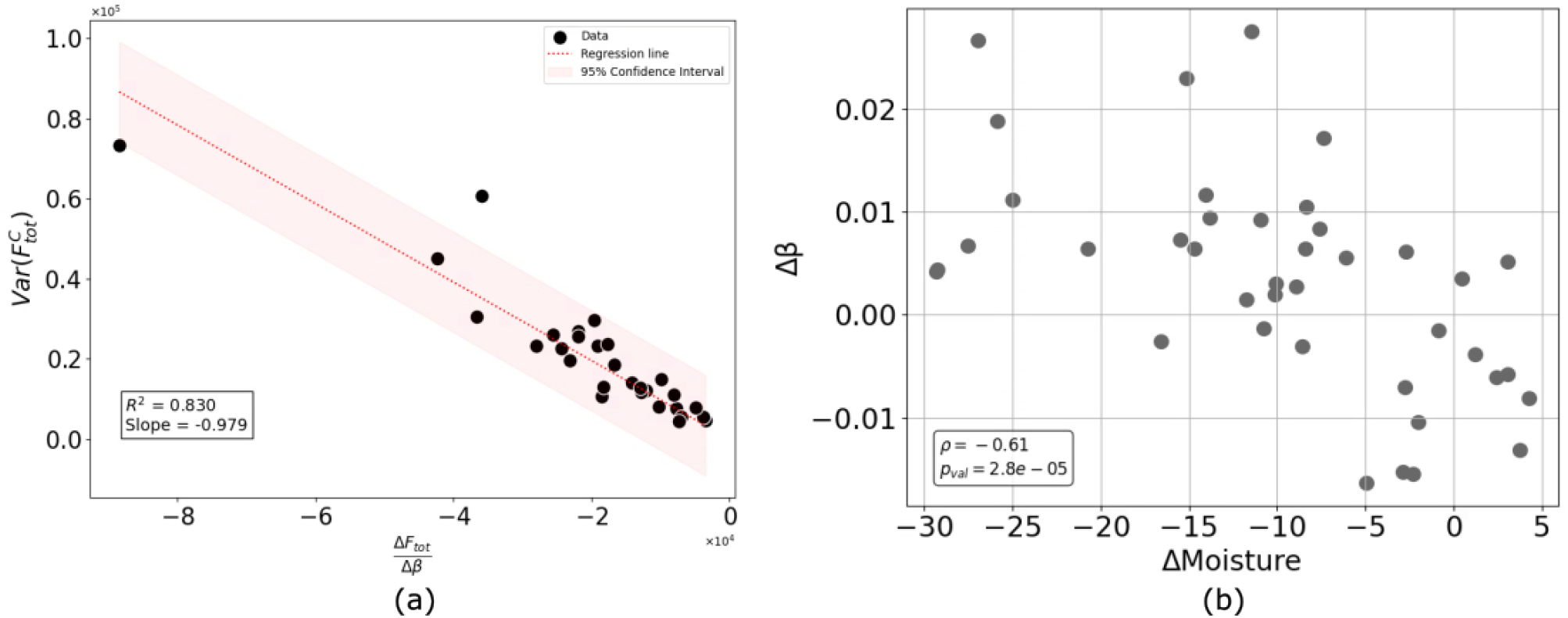
Fluctuation-Response Relation in an experimental soil food web. The relationship shows that the variance of the total flux at equilibrium negatively correlates with the amount of flux change caused by the perturbation (a). In panel b, the intensity of the perturbation Delta-Field moisture(difference between Control and Drought) is directly related to the variation in the structure of the network represented by the variation in the beta Langrage multiplier between control and the drought perturbation.

The Fluctuation Response Relation implies that a system characterized by large fluctuations at equilibrium, will respond to perturbations with a greater change than a system characterized by small fluctuations. The major implication is that the fluctuations at equilibrium are predictive of the variation following a perturbation. Our results show that an energetic description of food webs can be successfully integrated with a maximum entropy network description to bridge the gap between classic theory of food web stability and the experimental measurements of network stability properties. The step forward can be made by using maximum entropy to describe the fluctuations of the natural system around its typical configuration at equilibrium, which also links the fluctuations to the response of the network to perturbations. As energy fluxes can be experimentally measured, using the maximum entropy approach proposed here, it is possible to quantify them and so estimate the response to perturbation. To add to this, we offer a novel perspective on the relationship between the classic theory of food web stability and experimental network stability measures, as well as a new interpretation of the entropy metrics used in the maximum entropy approach. These metrics can be seen not only as abstract quantities but as meaningful indicators of network robustness and system variability. The same applies to any type of directed quantitative network where the link weights might be in different units, such as units of transferred information, and the quantitative fluxes between the nodes directly govern the dynamics of the node size. More broadly, the methodological framework developed here is adaptable, with the right caveats, to networks with varying structural constraints, making it suitable for investigating stability properties across a wide range of systems. Future studies could explore its application to networks subject to different forms of perturbation, with ensemble construction tailored to reflect the specific features of each system.

## II. METHODS

### A. The dataset

We analysed 60 replicates of the same soil food web with data collected from a large scale rain-out shelter experiment conducted from May to September 2016 in mesotrophic grasslands across three regions in south, central and north Britain. Details are given in [35]. The dataset consists of 30 control and 30 perturbed (i.e. drought) paired plots (a pair being made by a control plot and a perturbed plot). The rain-out shelters simulated a drought scenario equal to or more severe than a 100-year drought for these locations and details are given in [23]. The plots were also fenced to prevent grazing and other types of disturbances. After 60 days, drought shelters were removed and plots were sampled and assessed for soil organisms four times at shelter removal, and then 8, 21 and 60 days after the removal of the shelters. Details of the experiments and how the resulting food webs were measured are reported [23, 36]. We built the soil food webs with: i) the topology of the network (deterministic and static because data are aggregated at the high level of 13 major trophic groups whose links are known; see Figure 3), ii) the body mass of an individual (experimentally measured in mg), iii) the individual metabolic rate of an individual (measured in Joule per hour, and estimated using an allometric relationship that links measured body mass and environmental temperature to metabolism), and iv) the number of individuals in the population per unit of space.

#### Box II:Predicting responses to future perturbations from current food web fluctuations

**Setup**. Maximizing Shannon entropy *S*[*P*] = − ∑_*s*_ *P*(*G*) ln *P*(*G*) subject to the normalization constraint ∑_*s*_ *P*(*G*) = 1 and the expectation value ∑_*s*_ *P*(*G*)*C*(*G*) = ⟨*C*⟩, one obtains the exponential family

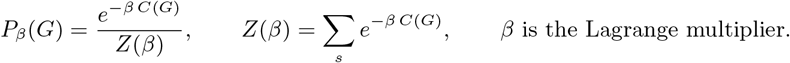

**Basic identities (cumulant generators)**. Considering ln *Z* (the free energy [34]), we can compute its second derivative with respect to the parameters.

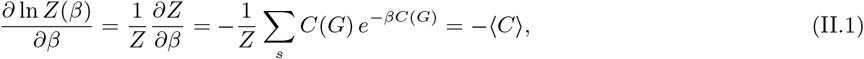

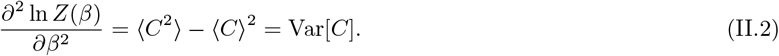

**Fluctuation–response relation**. Differentiating the first identity with respect to *β*:

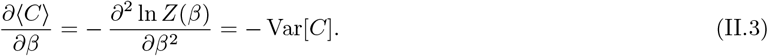

Writing this explicitly in terms of *Z*:

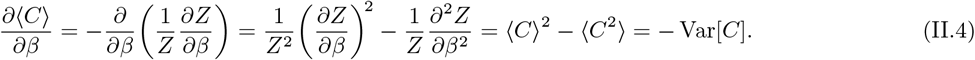

**Interpretation** The left-hand side of eq. (II.4) is the *response* (susceptibility) of the observable to a perturbation in the conjugate parameter *β*. The right-hand side is the *fluctuation* at equilibrium, Var[*C*] [24]. Thus, for a small perturbation Δ*β*:

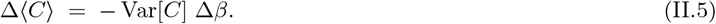

In our application, we set *C* ≡ *F*_tot_. Knowing the value of *F*_tot_ before the perturbation allows us to predict its post-perturbation change through its variance.

**FIG. 3:**
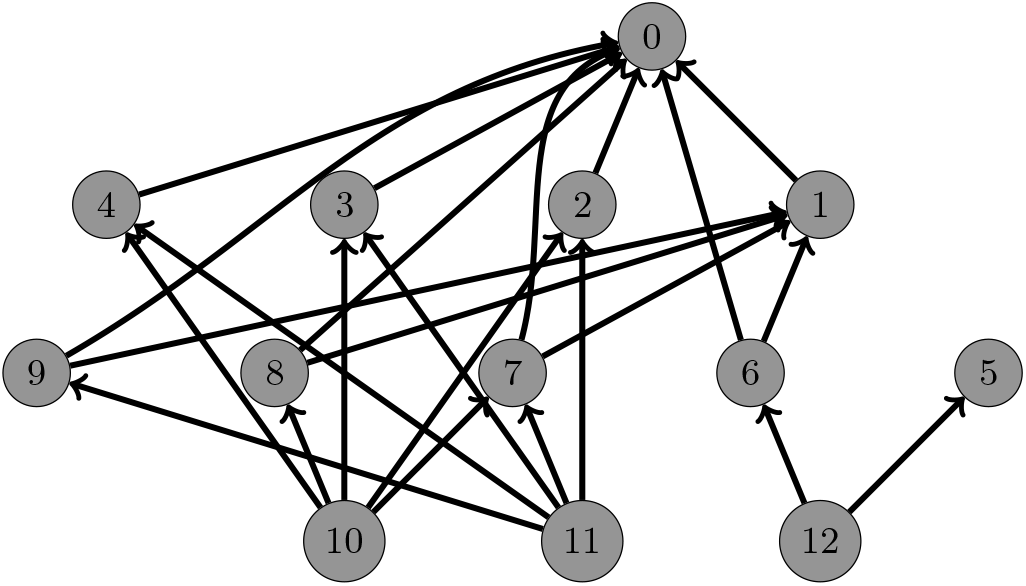
The topology of the analyzed food webs. Nodes ∈ {10, 11, 12} are the basal resources, while nodes ∈ {0, 5} are top predators. All other nodes represent consumers.

### B. Energy flux calculations and Population Dynamics

The amount of energy flowing from a resource species to a consumer species was measured in units of energy (in our case Joule) per unit area in the unit of time (one hour in our case), that is a flux. The energy is moved between species in the form of chemical energy “trapped” in the biomass consumed by the consumer. That biomass, once ingested, is partially (i.e. there is a loss) assimilated and metabolised to sustain the standing crop of the consumer, and its growth and reproduction. Also, the energy entering a trophic species, after accounting for the energy lost through natural processes (such as metabolism) should be equal to the energy exiting, leading to the mass balance equations (See Box.III).

Top predators use all the energy to maintain their biomass and as they give energy to no other species, while basal resources are assumed to receive energy only from the environment. This is important in the formalism of an energetic food web because a food web is a directed network with all links directed from the bottom to the top of the food web. That also means that in an explicit model the basal resources must either have a constant supply rate (external to the food web) or be modelled as a logistic population and so have an implicit resource with an equilibrium between resource supply and consumption. This point is important to achieve local stability dynamics [30, 33, 37].

To link the matrix **F** to an underlying population dynamics model, we use interconnected, non-linear differential equations based on the Lotka-Volterra type [2, 30], with a Type I functional response (Box III). The functional response can be changed but as we are introducing our framework for the first time, we limited ourselves to Type I responses. The set of equations used to describe the system are similar to those described in [30].

### C. Stability Metrics

The interaction strengths, *α*_*ij*_ are the elements of the Jacobian community matrix **J** of our system of differential equations, defined as the partial derivatives of the differential equations near equilibrium:

#### Box III: Mass-balance framework linking energy fluxes and population dynamics

**Step (a): Energy fluxes**. The energy flow from a resource species to a consumer species is measured as a flux (Joule per unit area per unit time). Energy enters consumers in the form of biomass ingested, part of which is assimilated and metabolized to sustain biomass, growth, and reproduction. For each species *i*, we define incoming and outgoing fluxes:

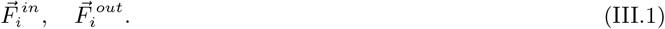

Mass-balance requires that energy entering equals energy exiting once metabolic losses are accounted for.

**Step (b): Mass-balance equations**. At equilibrium, the fluxes satisfy

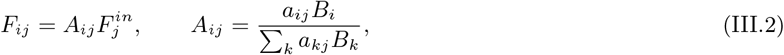

with *a*_*ij*_ ∈ {0, 1} defining the network topology. This leads to the balance equation

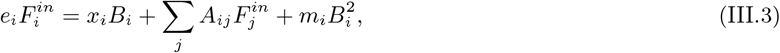

By knowing 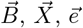, and 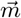, it is possible to derive the matrix of fluxes *F* = {*F*_*ij*_}.

**Step (c): Linking to population dynamics**.

The flux matrix is connected to Lotka–Volterra differential equations with a Type I functional response following [30]:

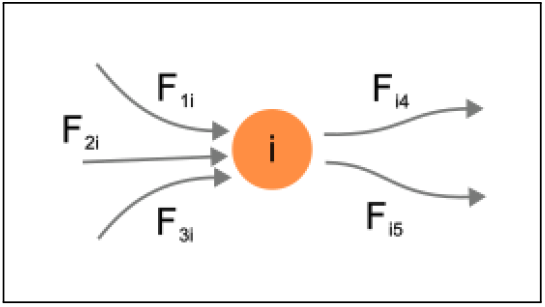

for basal resources,

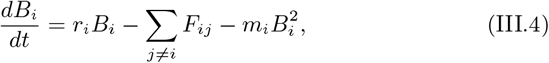

and for consumers,

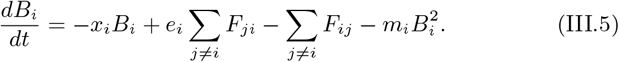

Incoming and outgoing energy fluxes in the mass-balance framework.

Here *r*_*i*_ is the intrinsic growth rate of basal resources, *x*_*i*_ the metabolic rate, *e*_*i*_ the assimilation efficiency, *m*_*i*_ the self-limitation term, and *F*_*ij*_ the energy flux from *i* to *j*.

**Step (d): Canonical ensemble and randomization**. The population dynamics defined above can be sampled within a canonical ensemble, where randomized flux configurations are generated consistently with the mass-balance constraints. This allows comparison between the observed dynamics and randomized counterparts.

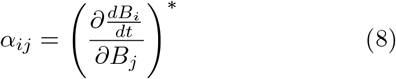

The derivation, based on the linearization of the system of differential equations using a first-order Taylor series approximation, is very well known [1]. The knowledge of the energy fluxes allow parametrising the Jacobian of the corresponding population dynamic model (See Supporting Information Sec. 5 for further details).

Stability is a central topic in the theoretical ecological literature and the term stability has been associated with different meanings and so measurements [9]. Here we refer to two different types of metrics. First, the empirical ones, which are independent of descriptive models, relying solely on empirical data gathered from the field. Among those metrics, there is the coefficient of variation [38]:

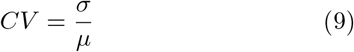

that is the ratio of the empirical standard deviation (*σ*) to the empirical mean (*µ*), offering insight into the extent of variability relative to the average state of a particular system. Specifically, in the context of population ecology, the coefficient of variation becomes instrumental in assessing the spatial or temporal variance in population density (either of individual species or the total community as the sum of the individual species). Second, there are the model based metrics (See Supporting Information Sec. 4). Among those, the neighbourhood or local stability received most of ecologist attention in the quest for stability criteria [2, 33]. Local stability describes the behaviour of a system in the proximity of its equilibrium point.

The stability of the equilibrium is evaluated with the eigenvalues *λ*_*i*_ of the Jacobian matrix [1, 2]: if the maximum among all the real parts of the eigenvalues is negative, the perturbations *x*_*i*_(*t*) decay to 0 (with or without oscillation, depending on whether the eigenvalues have non-zero imaginary parts). Since the eigenvalues are related to the rate of perturbation decay, they can also be used to quantify the resilience of the system, that is the rates at which the system recovers from a perturbation, formally quantified as the inverse of the Return Time (RT):

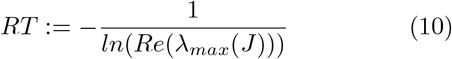

Where *λ*_max_(*J*) is the least negative eigenvalue. Great resilience corresponds to enhanced stability. Vice versa, diminished resilience (or longer return time, *RT*) means a prolonged time for the system recovery. That increases the likelihood of more perturbations catching the system during the recovery phase [33].

The life cycle of the species under investigation provides a rough estimate of the scale of return time, which is useful when parameterising the models as per Supporting Information Sec.(5). Another metrics we considered and that is directly computed from the Jacobian matrix is the *Reactivity* [39]. Mathematically, reactivity is measured as the leading eigenvalue of the symmetric part of the Hermitian of the Jacobian matrix. Reactivity identifies the occurrence of transient growth following a perturbation, preceding the return of a system with a stable equilibrium to its initial state. If a system is locally stabile, positive values of Reactivity means that there will be an initial amplification of the perturbation immediately after the perturbation hits, followed by decay. If negative, the perturbation will just exponentially decay back to the equilibrium. Reactivity therefore captures the short term response to the perturbation. We also derived the Lyapunov function to identify criteria of global stability for our food web (Supporting Information 7).

### D Network model, fluctuations and dynamics

We estimated the parameters 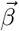 and the Bolzman distribution, that is in this case the probability of observing a matrix **F** given the topology of the food web (*a*_*ij*_) and the Lagrange multipliers 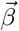 in graph Hamiltonian H 1, following the procedure detailed in the Supplementary Information 1.

In principle, however, the model can in the future be extended to fluctuating topologies, but subjected to certain constraints to forbid impossible links (for example, there can be no directed link from a top predator to a consumer or to a basal resource because, obviously, consumers and basal resources do not eat top predators).

The Lagrange multipliers 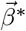 that correspond to the observed constraints, 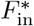 and 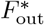, are found by the maximum likelihood method [22, 27]. With those at hand, we can compute the numerical value of the entropy (See Box I Eq. I.5).

The canonical ensemble generated from the observed matrix of fluxes *F* ^∗^ can be sampled numerically to generate a desired number of randomised flux matrices that on average respect the constraints 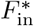 and 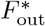 for each node *i*. As each sampled matrix can be interpreted as fluctuation around the equilibrium, the Jacobian can be recalculated for each sampled flux matrix. The equilibrium values of *B*_*i*_ must be recalculated as well in order to ensure that the new, randomised system is at equilibrium and feasible, with all species having positive densities.

Practically, from the ensemble, we generate a matrix that we indicate as *F*^samp^ and from it calculate 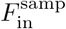 and 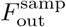. At this point,we search for the values of 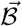 such that:

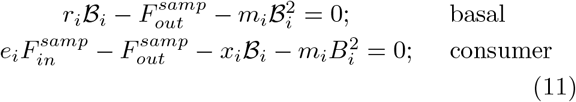

The vector 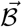 lists the biomass values that realize the new equilibrium state of the system, which can be assessed using stability metrics such as the leading eigenvalue. The computer code and an example dataset used to perform all computations are available at: https://github.com/VirginioClemente/Energetic-Food-Webs.

## AUTHOR CONTRIBUTION STATEMENT

GVC, DG and TC developed the concept and theory of the paper; GVC leaded on data analysis with TC and DG support; GVC and TC leaded on the writing with contributions from all co-authors; TC, CM, JM, RB, DJ, FdV and ME provided the food web data; All authors intellectually contributed to the interpretation and discussion of results.

## ACKNOWLEDGMENT

This work is supported by the European Union - NextGenerationEU - National Recovery and Resilience Plan (Piano Nazionale di Ripresa e Resilienza, PNRR), project ‘SoBigData.it - Strengthening the Italian RI for Social Mining and Big Data Analytics’ - Grant IR0000013 (n. 3264, 28/12/2021) (https://pnrr.sobigdata.it/). This publication is part of the projects “Network renormalization: from theoretical physics to the resilience of societies” with file number NWA.1418.24.029 of the research programme NWA L3 - Innovative projects within routes 2024, which is (partly) financed by the Dutch Research Council (NWO) under the grant https://doi.org/10.61686/AOIJP05368 and “Redefining renormalization for complex networks” with file number OCENW.M.24.039 of the research programme Open Competition Domain Science Package 24-1, which is (partly) financed by the Dutch Research Council (NWO) under the grant https://doi.org/10.61686/PBSEC42210. This study was supported by the Natural Environment Research Council (NERC) Soil Security Programme grant initiated and led by RDB, with component grants NE/M017028/1 to R.D.B. and F.d.V., NE/M01701X/2 to D.J. and E.M.B., and NE/M017036/1 to T.C. and M.E. We are very grateful to the landowners and farmers for allowing us to perform our experiment and sample their fields. RDB was also supported by the European Research Council (ERC) SoilResist (Grant agreement No. 883621), and FTdV was also supported by a Starting Grant SHIFT-FEEDBACK (Grant agreement No. 851678) from the European Research Council. TC was also supported by Research Ireland (20/FFP-P/8584). GVC acknowledges the JRC Exploratory Research Programme, which supported the Exploratory Research Project “REloading the Anthropocene to Decode Mass Extinction (README)”.

## SUPPORTING INFORMATION

### 1. The Conditional Reconstruction Method

Here, we summarise the Conditional Reconstruction Method (CReM) proposed by Parisi et al. [29], which separate the estimation of parameters governing topology from that of the parameters controlling the weights of the link. The model estimates the probability of observing a weighted graph W, Q(W), which can be expressed as the product of the probability of observing a topology A, P(A), and the probability of observing a weighted graph W given the topology A, *Q*(*W* |*A*).

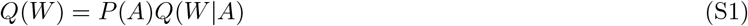

We focus on the estimation of the probability *Q*(*W*|*A*), assuming P(A) as given. The objective of the CReM is to obtain the distribution of weighted configurations given prior information on the underlying adjacency, binary matrices. The methods search for a functional form of *Q*(*W*|*A*) that satisfies *Q*(*W*|*A*) = 0 for *W ∉ W*_*A*_ (where *W*_*A*_ is the set of possible weighted configurations for a given topology **A**), while maximizing the conditional entropy:

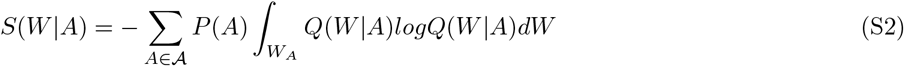

subject to the following constraints

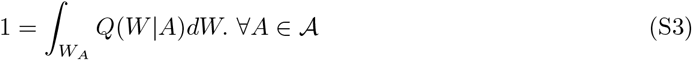

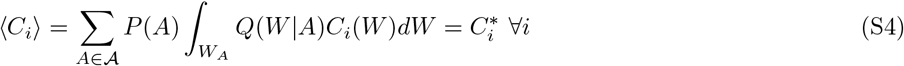

The maximization procedure is carried by employing the method of the Lagrange multipliers. So after writing the Lagrangian:

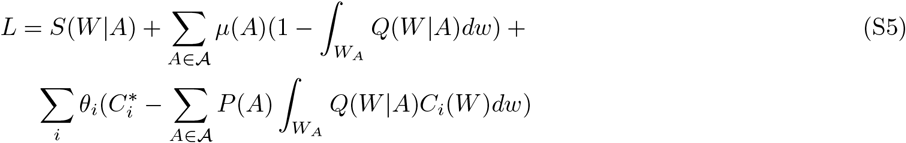

We can differentiate w.r.t. *Q*(*W* |*A*) and find the expression of *Q*(*W* |*A*) that would be the form of ERGM [21]:

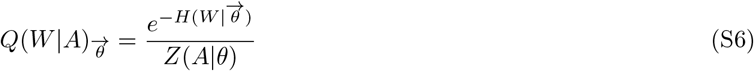

The dependency of the function on the set of Lagrange multipliers, denoted as 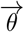, is explicitly evident. These multipliers serve as free parameters that require determination to precisely mirror the actual values of the observed constraints, 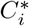. In addressing this issue, the authors in [29] introduced an extended form of the likelihood, termed the “Generalized Likelihood”:

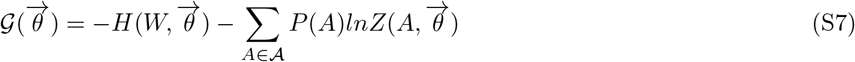

The optimal set of parameters, represented by 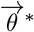, is the one that maximize the likelihood. In our application of this method, we can also derive the relation that links the conditional entropy, defined in eq. (S2), with the generalized likelihood, both computed in 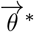.

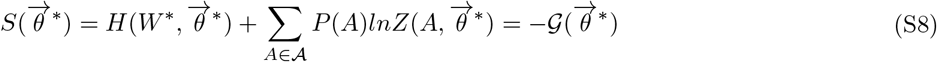

This allows us to derive the entropy value used in the main paper from the generalized likelihood, connecting it with other quantities of interest analyzed.

For our specific application, we constrain the expected values of the in- and out-strengths (*s*^*in*^, *s*^*out*^), which are the energy fluxes, given a fixed, known topology. Defining *w*_*ij*_ as the *i, j* element of the weighted graph *W*, we can express the strengths for each node as:

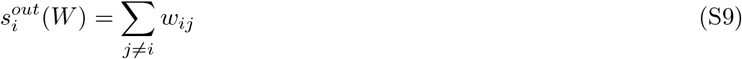

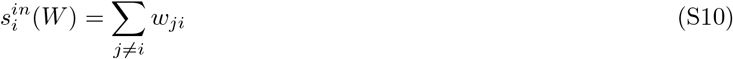

The graph Hamiltonian then is:

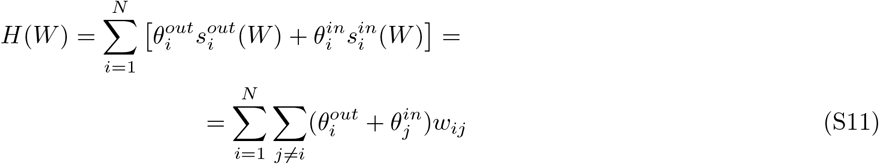

That can be associated with the following partition function:

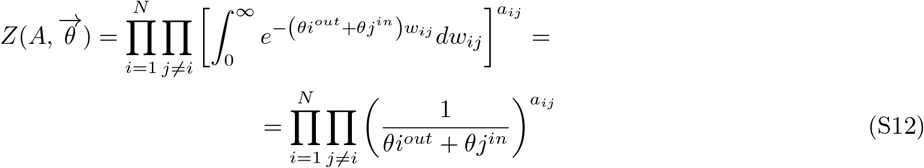

And using equation (S6):

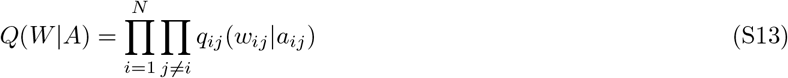

with:

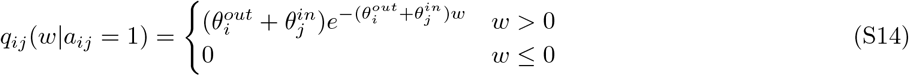

The generalized likelihood associated with this system has the following expression:

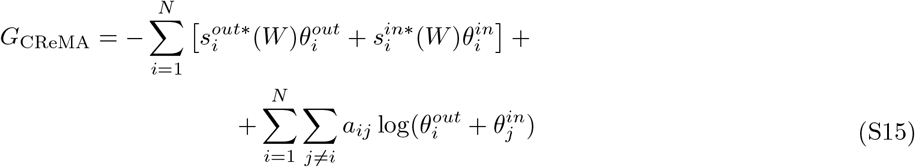

The procedure used to find the value of the parameters 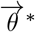 requires maximizing the generalized likelihood (Eq. S15) with respect to these parameters, which is equivalent to solving the following system of equations:

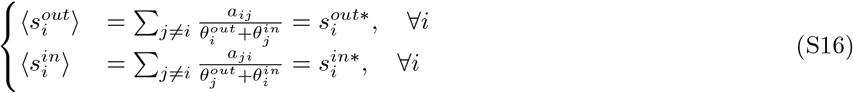

For further details on the calculations, see Parisi et al.[29].

Once the parameters are estimated, these can be used in eq. (S13) to sample a graph W and to find the numerical value of the entropy associated with the described system.

The form of the model also allows examining the contribution of each (weighted) link to the overall value of the entropy. By construction, the link weights between each node, which is the quantity of energy flowing between two nodes, is:

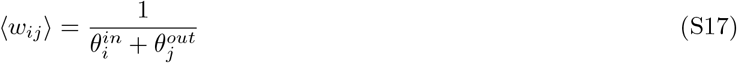

This estimate coincides precisely with the value calculated through eq. (III.2) of the Box III for each food web. The implication is that the least biased way to allocate energetic contributions to every single from other nodes is to do it proportionally to the biomasses of the resource nodes. Also, the sequence of the steps we employed to estimate the Maximum Entropy network ensemble allows the use of the Mass Balance equation to compute fluxes, a step essential for constructing the complete structure. This result has significant implications in assessing the contribution of the weight of every single link in computing the value of the entropy. Indeed, by combining eq. (I.5) with eq. (S17), it can be observed how the contribution to the system’s entropy, and hence, as we will demonstrate in the main text, to its stability, is proportional to the weight of the links. Indeed, one of the components in the entropy formula, specifically the logarithm of the partition function, is given by:

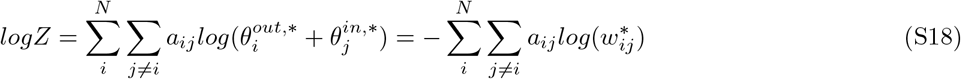

If we place it back in the expression of the entropy, we find that, in our framework, the stronger links are those that affect most significantly to the system’s entropy.

### 2. Total flux as macro-constraint

In this section, we construct a maximum entropy model in which the total flow of the system is constrained simultaneously with the topology. The Hamiltonian associated with becomes:

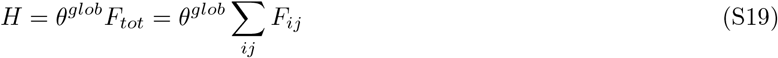

The resulting model, in this case, implicitly redistributes the weight evenly across all available links in the network, resulting in links that will all have the same expected weight. We solve the model using a variation of a specific instance of the CReM, denoted as CReMb [29]. In our scenario, assuming 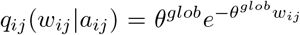for *w_ij_* > 0 and 0 otherwise, starting from the Hamiltonian in eq. (S19), the generalized likelihood takes the following form:

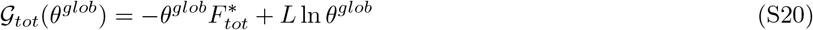

Where the asterisk indicates the measured value. Deriving with respect to *θ*^*glob*^, we obtain:

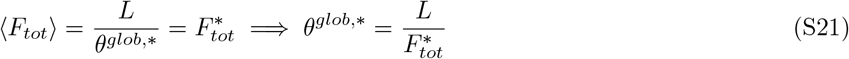

This implies that the conditional entropy (eq. S8) becomes:

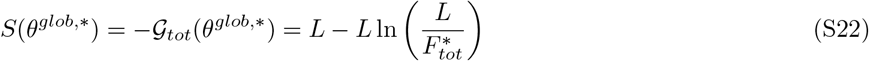

The entropy increases when the total flux in the system increases. Beyond the expression for entropy, once the *θ*^*glob*,∗^ parameter is obtained, we can proceed to calculate the variance associated with the total flow, which will be equal to:

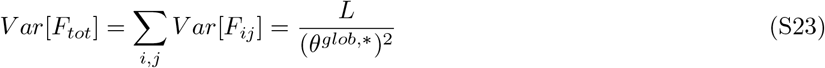

### 3. Hybrid Model

We introduce a hybrid model to connect local and global constraints in our application of the Fluctuation–Response Relation. Compared with CReMa (Sec. 1), it is obtained by the reparametrization 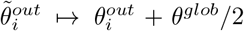 and 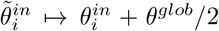, which directly links the local formulation to the model constrained only by total flux. The corresponding Hamiltonian is

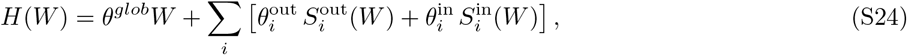

which can be rewritten as

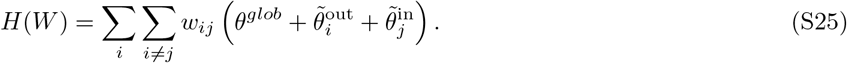

The corresponding partition function is

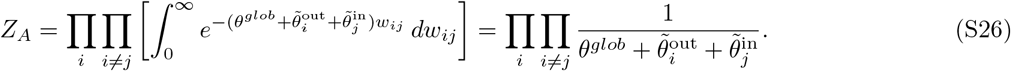

This yields the probability distribution

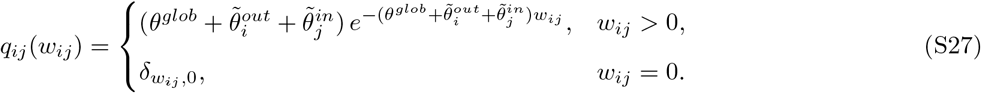

Accordingly, the generalized log-likelihood becomes

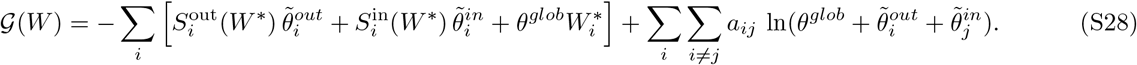

From this point, we can estimate the values of the Lagrange multipliers by following the same procedure described in Sec. 1, i.e. by maximizing the log-likelihood. This procedure corresponds to solving the following set of equations:

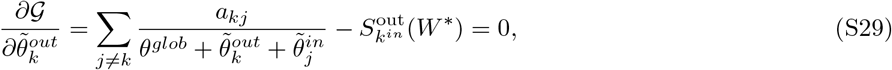

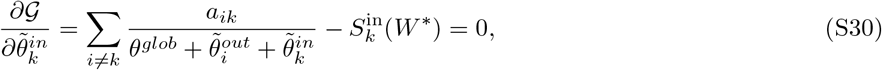

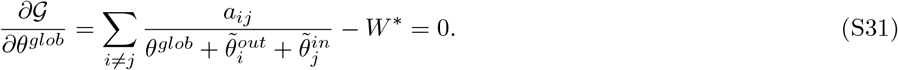

In order to derive the relation used for the Fluctuation Response Relation, we focus on the second derivative of *G* with respect to the global parameter *θ*^*glob*^, for which we want to obtain this relation. By computing this derivative, we find:

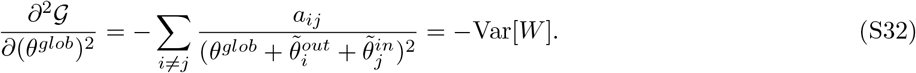

Since *W* ^∗^ is nothing more than the sum of the *w*_*ij*_, which are already determined by the *S*^in^ and *S*^out^ constraints, in this context there is a redundancy that allows us to choose the global parameter *θ*^*glob*^ independently.

Therefore, while the variance of *W* can be obtained from the CreMa by using the variances of the individual *w*_*ij*_ and exploiting the relation Var[*W*] = ∑_*i,j*_ Var[*w*_*ij*_],

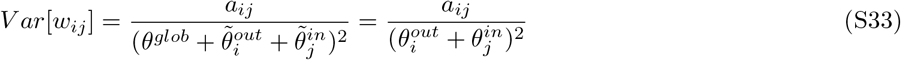

the global parameter *θ*^glob^ can be chosen so as to satisfy the global constraint. Specifically, we set *θ*^glob^ equal to the value that solves the CreM model with the global constraint defined in Sec. 2:

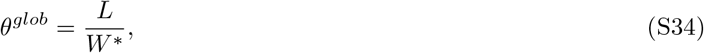

where *W* ^∗^ is the empirical total weight and *L* the number of links.

### 4. Stability Metrics

From the Jacobian matrix it is possible to derive several stability metrics. Let us consider a system of differentiable equations with fixed parameters:

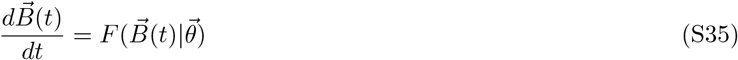

Where 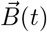 is a random vector, 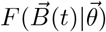 represents a function applied to the vector 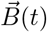, and 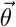 is the vector of fixed parameters on which the function depends. We assume that there exist at least one equilibrium point 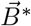, and we study the dynamics around this point by taking a small perturbation for each element *i* of this vector:

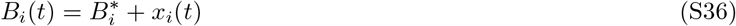

Employing a Taylor expansion allows us to linearize the system, thereby facilitating the examination of the perturbations’ evolution:

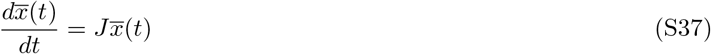

Within this linearized system, *J* represents the Jacobian matrix, with each element *α*_*ij*_ defined as follows:

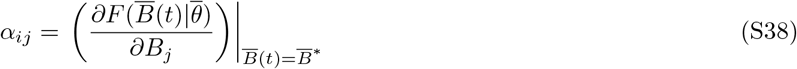

This means that given our data, we can use our estimate of fluxes and interaction strengths from our dynamic system in eqs III.4 and III.5 to derive eq. S38. Another metrics we considered is:

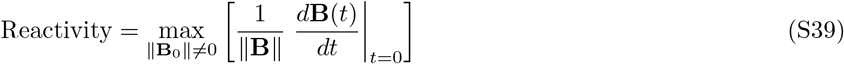

Where **B**_0_ is an initial perturbation. This has been shown [39] to be equivalent to:

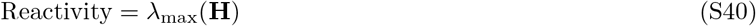

Where **H** = (**J** + **J**^*T*^)/2. Reactivity is measured as the maximum eigenvalue of the symmetric part of the Hermitian of the Jacobian matrix.

### 5. Jacobian Parameterization

The interaction strengths, *α*_*ij*_ are related to the fluxes by:

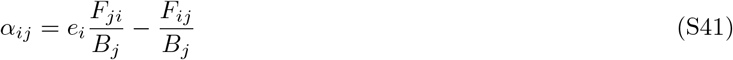

With additional simplifications applied to the eq. (S41), it becomes straightforward to demonstrate that, by considering the values of the flux matrix **F**, the impact of consumers *j* on resources *i* can be simplified in the following expression 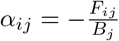. Similarly, the influence of resources *i* on consumers *j* can be written as 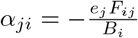. Moving to the diagonal elements of the Jacobian, we get:

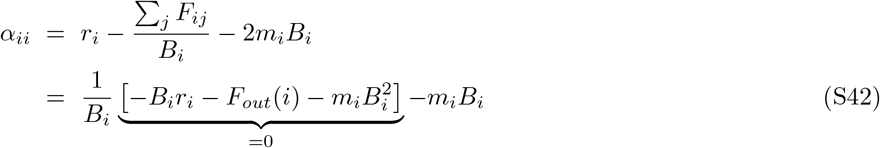

For basal resources, while for consumers:

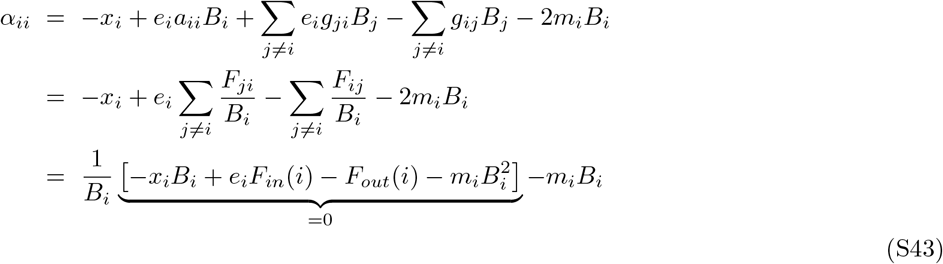

It can be noted that in both eqs.(S42) and (S43) the terms written between squares parenthesis are exactly the expressions of the mass-balance equations (2) and (3). Hence these terms are by definition equal to 0. This consideration implies that all the diagonal terms of the Jacobian matrix can be written as:

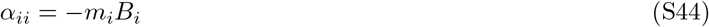

Given the unknown nature of the term *m*_*i*_, we have to propose a proxy for its value. In doing so, we follow the approach adopted by [30], where the term *m*_*i*_ is assumed to be proportional to the death rate of each specie.

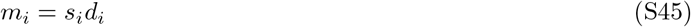

Where *d*_*i*_ is the death rate and *s*_*i*_ is the proportional factor.

How to select *s*_*i*_? We assume that, being the observed system near or at some local equilibrium (after all, it was observed and measured over a period of time of several months and the system is still there!) the maximum among the real parts of the eigenvalues of the Jacobian matrix must be negative. The return time should then be in the order of the growth rates of the basal species under investigation. In our case, we use the dominant plant species of the grassland in the system, with return rates of a few years max (based on the average population intrinsic growth rate). There are no other practical solutions unless one could measure return time experimentally, which is challenging but in principle possible. The need to work on the basis of assumptions around mi and si arises from the logistics form of the equation, but it’s essential to ensure that the population model can have some feasible and non-trivial local equilibrium. This is a well-known theoretical point [1, 2, 33].

### 6. Partitioning the entropy contributions

One might wonder if the observed relationship between entropy and other metrics is merely a consequence of a dependence of entropy on total flow, suggesting that entropy doesn’t convey any additional information beyond this quantity. However, entropy not only encompasses information about the total energy flow through the system but also reflects how this energy is distributed among its various components. Operationally, to disentangle the factors contributing to the entropy, we’ve adopted a methodology that allows us to standardize the flows within each individual food web. By introducing a multiplicative factor *α*, we ensure that each food web has the same total flow by scaling each energy channel by this factor. As a result, the only variation between different systems will be how this energy is distributed. However, before proceeding with the calculation of this new metric and its relationship with other properties, it’s crucial to show and emphasize that this transformation does not affect the system’s population dynamics, thereby leaving the Jacobian unchanged. This means that any observed relationships will be genuine, without alterations due to data transformation. First, if we simply multiply all the biomasses for a generic constant *α*, by substituting the biomass values in Eqs. 2 and 3, with the same biomass multiplied by *α*, we obtain teh following equations:

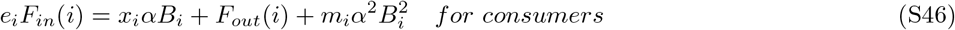

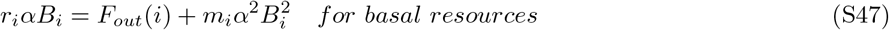

Thus, the modifications to the flows induce the following changes on the elements of the Jacobian:

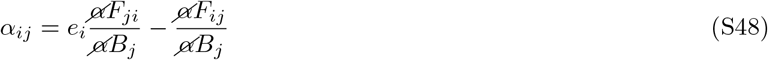

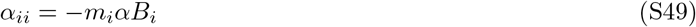

Following the procedure initially defined to search for *m*_*i*_, in order to maintain the system’s return time (and thus the eigenvalue), we would have a value of 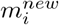 that is equal to 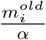. This implies that the Jacobian describing the system does not change.

Instead, we get a change in the fluxes flowing in each connection. Those changes can be resumed in the following equations:

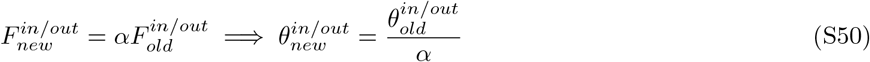

This induces a change in the expression of the entropy that is directly dependent on the value of *α*, described by to the following expression:

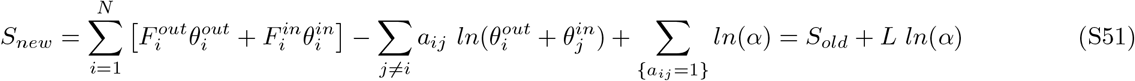

We obtain that the new entropy will be equal to the old one plus a constant, depending on the value of *α*. This implies that if we increase the tot flux in the system, keeping the initial distribution, we can see a change in the entropy that corresponds to *L ln*(*α*). The *α* can be set based on the highest total flux observed across the food webs in the dataset. Each food web’s flux can be normalised by dividing it by this maximum value, ensuring that *α* consistently lies between 0 and 1. Thus, by reinterpreting *α* as a parameter useful for standardizing flows, it is possible to show that the value of *L* ln(*α*) is a linear transformation of the entropy calculated from a model where the total constraint is the total flow (See SI. 2).

This means that in regression exercise to relate entropy to other metrics, it is possible to use 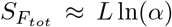. Standardizing the flows allow to view *S*_*new*_ as the contribution of entropy arising solely from the flux distribution. For example, we examined the correlation between the two entropies and CV, ℜ(*λ*_max_), and Reactivity. For a comparative assessment of models with multiple variables against the one solely based on entropy, we used the Bayesian Information Criterion (BIC). We renamed *S*_new_ as *S*_het_, and *S*_old_ (the entropy of the system where both characteristics are considered) as *S*_Full_. Results (Table I) show that *S*_het_ is always required as a linear predictor of all three stability metrics, and in the case of CV it is in fact the only predictor that survives model selection. For reference, the model using *S*_Full_ alone yielded BIC values of 165 for CV, 155.4 for ℜ(*λ*_max_), and 167 for Reactivity. The main takeaway is that *S*_Full_ captures both contributions in a more parsimonious way. We note that the observed relationships are non-linear, but we present the comparison in this form to make it more intuitive and to emphasize a theoretically grounded result.

**TABLE 1:**
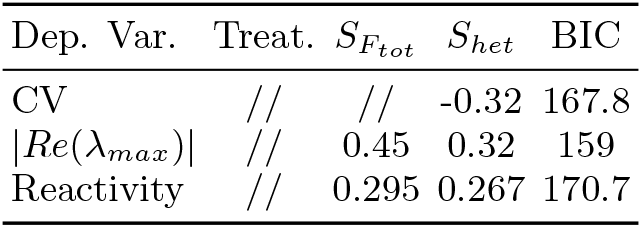
Coefficients and BIC for the regression models with *S*_*tot*_ and *S*_*het*_, and treatment. // indicates that the coefficient associated with the column variable is not significant.

### 7. Lyapunov Function and global equilibrium points

This section will detail how to derive the conditions for verifying global stability at the equilibrium point under consideration. Our methodology involves formulating a Lyapunov function tailored to the analyzed system. Over the years, the global stability of Lotka-Volterra systems has been extensively studied and researched [40]. Goh was the pioneer [41, 42] in exploring sufficient conditions that guarantee the function, as represented by the equation

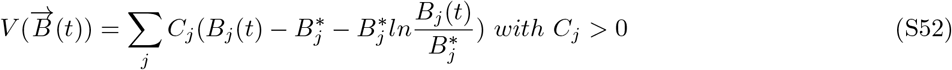

to be a valid Lyapunov function for the type of system we are currently examining.

Therefore, we investigate the behavior of this function within our specific Lotka-Volterra model and revisit the necessary steps to achieve the condition for global stability. It can be easily verified that 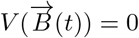 for *B*(*t*) = *B*^∗^, and that it is ≥ 0 for all 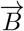. Therefore, the final step to verify the global stability of the equilibrium point involves calculating the time derivative of *V*:

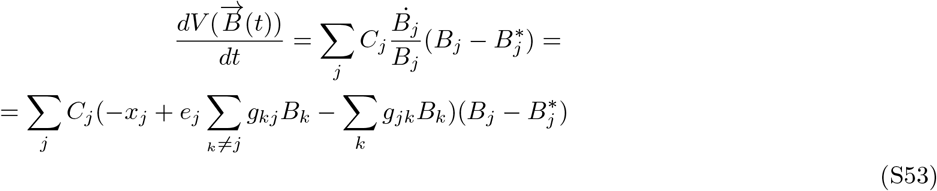

with *g*_*jj*_ = *m*_*j*_.

In equation (S53), we apply the equation for the consumers, as outlined in the following:

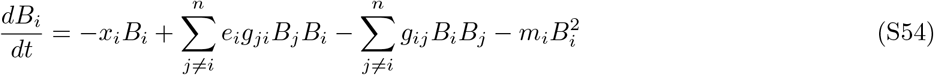

In the main text, following [30], we use 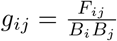. The derivation process for the basic resources follows a similar pattern. To determine whether our point represents a global equilibrium for the system, it is necessary to verify that:

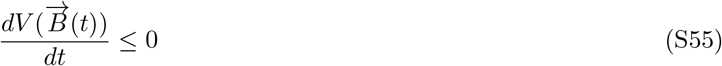

Now, by applying the conditions for equilibrium:

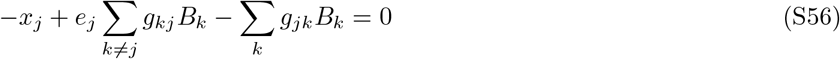

we get:

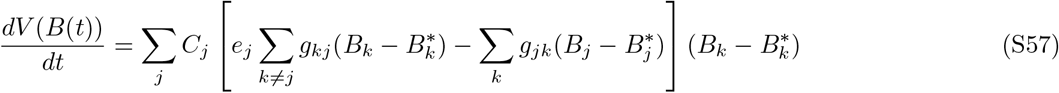

We can define the finite perturbation vector y with components 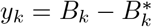, and then:

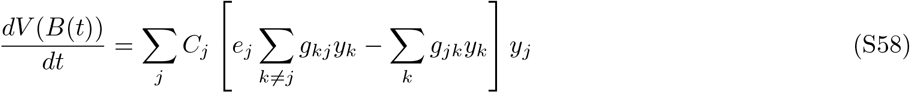

If we separate the sums, and relabel the second term:

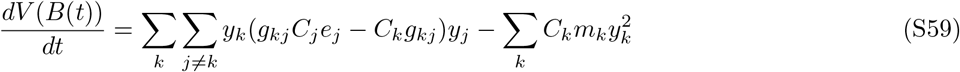

This is a quadratic form. To satisfy the condition in eq.(S55), our aim is to identify values for *C*_*j*_ *>* 0 for *j* ∈ 1 · · · *N* such that eq. (S59) is negative semi-definite. Utilizing the actual values from the adjacency matrix we are examining, we can explore a set of values for 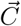. The goal is to ensure that all eigenvalues of matrix (4) are negative, which would validate the conditions in (S55). An analysis of the function with real-world data shows that it is always possible to find a 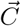 that meets these conditions. This leads to the conclusion that the equilibrium points 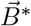 in our system are globally stable.

**FIG. 4:**
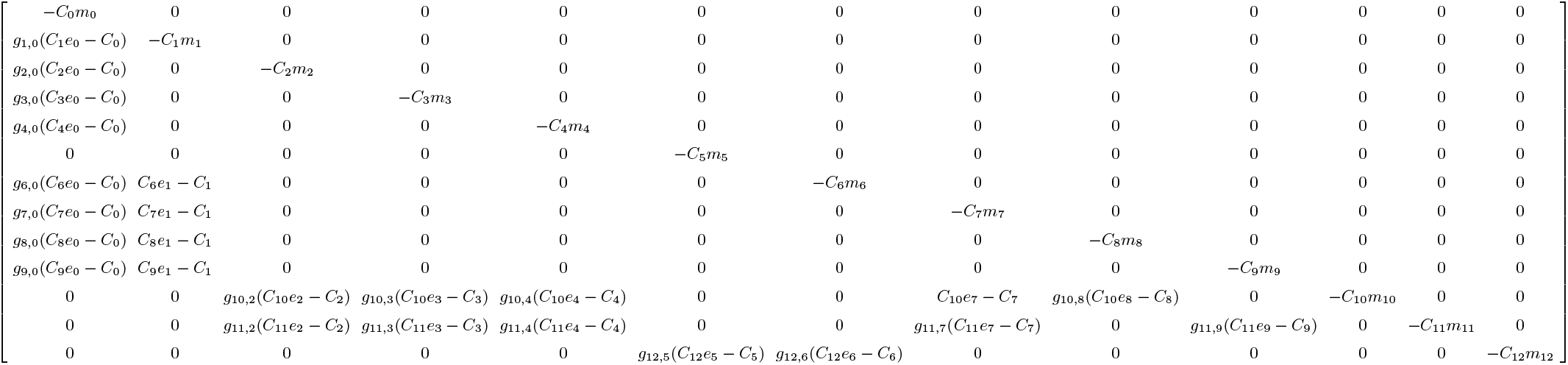
Matrix representing the elements of the operator associated with eq.S59.

### 8. Comparison Between Observed and Predicted Stability Metrics

**FIG. 5:**
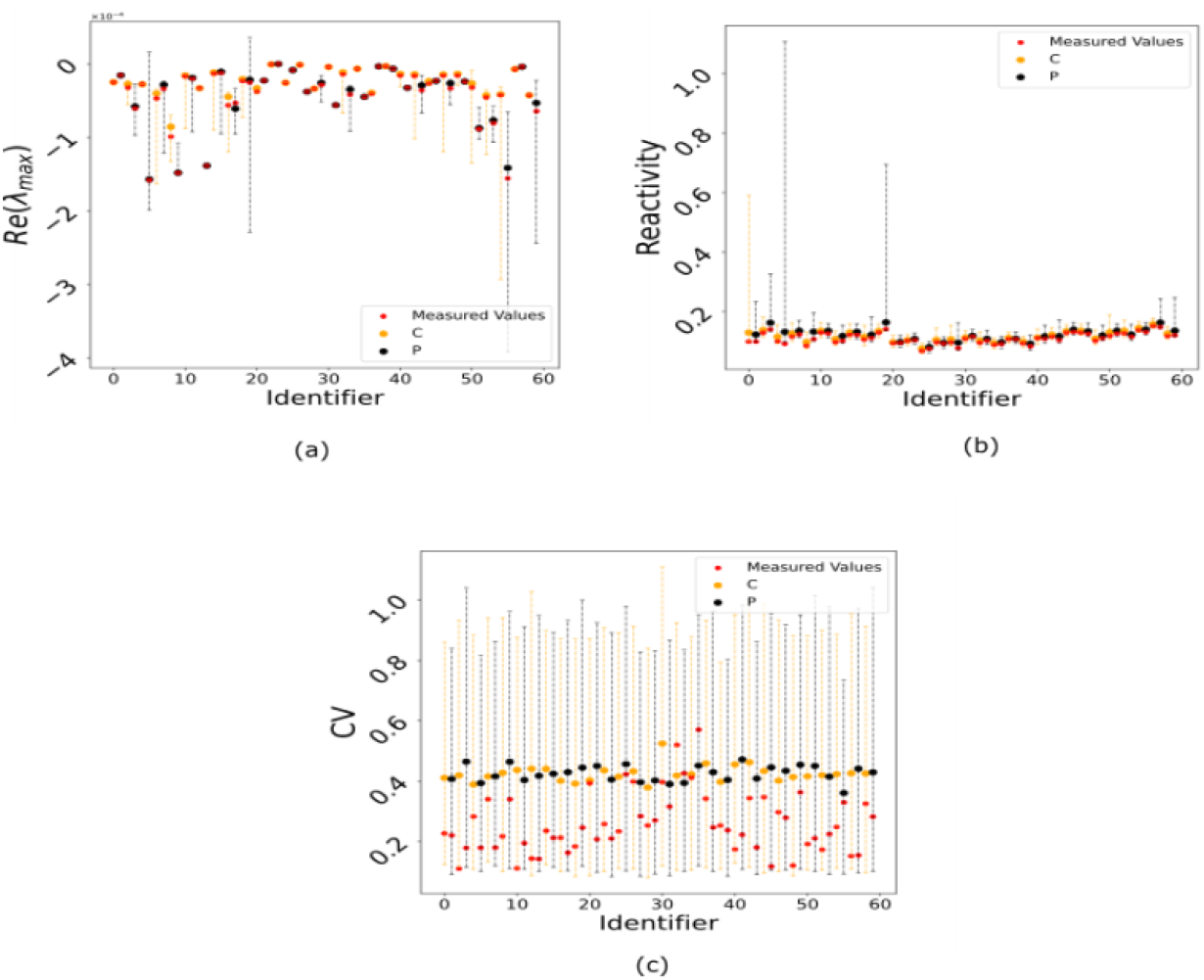
Comparison of Observed and Predicted Values in Food Web Dynamics. Observed values are denoted by red dots, and the ensemble medians, based on 1,000 simulations, are shown as blue dots. For figures (a) and (c), the vertical bars indicate the range from the 1st to the 99th percentile of the distribution. In contrast, figure (b) focuses on a narrower range, from the 20th to the 80th percentile.

## Notes

### Competing Interest Statement

The authors have declared no competing interest.

